## Supplementary information for "Trade-off between reducing mutational accumulation and increasing commitment to differentiation determines tissue organization"

### Supplementary information for Trade-off between reducing mutational load and increasing commitment to differentiation determines tissue organization

Márton Demeter
*MTA-ELTE "Lendület" Evolutionary Genomics Research Group, Pázmány P. stny. 1A, H-1117 Budapest, Hungary and*
*Department of Biological Physics, Eötvös University, Pázmány P. stny. 1A, H-1117 Budapest, Hungary*

Imre Derényi*
*Department of Biological Physics, Eötvös University, Pázmány P. stny. 1A, H-1117 Budapest, Hungary and*
*MTA-ELTE Statistical and Biological Physics Research Group, Pázmány P. stny. 1A, H-1117 Budapest, Hungary*

Gergely J. Szöllősi†
*MTA-ELTE "Lendület" Evolutionary Genomics Research Group, Pázmány P. stny. 1A, H-1117 Budapest, Hungary*
*Department of Biological Physics, Eötvös University, Pázmány P. stny. 1A, H-1117 Budapest, Hungary and*
*Evolutionary Systems Research Group, Centre for Ecological Research, Klebelsberg Kunó u. 3, H-8237 Tihany, Hungary*

*
†

#### A. Table of parameters

| Parameter name | symbol | description |
|---|---|---|
| Total number of cells generated | $N$ | Total number of cells generated during the expected lifespan of the tissue. |
| Number of hierarchical levels | $n+1$ | Number of hierarchical levels. |
| Number of cells at homeostasis at level $k$ | $N_k$ | Number of cells at level $k$ under homeostasis satisfied. |
| Differentiation rates | $\delta_k$ | Rate at which level $k$ produces differentiated cells under homeostasis. |
| Amplification factor | $\gamma_k$ | $\gamma_k = \delta_k/\delta_{k-1}$ describes<br>the ratio between the differentiation rates of neighboring levels rates. |
| Per cell rates of microscopic events | $r_k^{\circ\circ}, r_k^{\uparrow\circ}, r_k^{\uparrow\uparrow}, r_k^{\times}$ | Symmetric differentiation ($r_k^{\uparrow\uparrow}$),<br>asymmetric differentiation ($r_k^{\uparrow\circ}$ ),<br>symmetric cell division ($r_k^{\circ\circ}$),<br>and cell death ($r_k^{\times}$ )<br>for each level $0 < k < n$<br>and the division rate $r_0 = r_0^{\uparrow\uparrow}$ of the stem cells. |
| Number of driver mutations | $d$ | The number of driver mutations in cells. |
| Mutation strength | $s$ | Proportionality factor for driver mutations with which certain divisional rates are modified as specified in Eq. (10). |
| Driver mutation rate | $\mu$ | Probability of a mutation in each descendant cell following a division event. |
| Lifetime of the tissue | $t_{\text{life}}$ | The total lifespan of the tissue. |

TABLE S1. Table of parameters

#### Validating equation (23).

To validate equation (23), we performed explicit numerical simulations: We consider a hierarchical tissue with $n+1$ levels with $N_k$ wild type cell cells at each level at homeostasis and a uniform amplification factor ($\gamma_k = \gamma$) and corresponding rates of the four microscopic events: symmetric differentiation ($r_k^{\uparrow\uparrow}$), asymmetric differentiation ($r_k^{\circ\uparrow}$), symmetric cell division ($r_k^{\circ\circ}$), and cell death ($r_k^{\times}$). During the simulation we keep track of the number of cells with a given genotype, specified by the number of driver mutations, i.e., on level $k$ the number of cells with $m$ driver mutations at time $t$ is $N_{k,m}(t)$. We simulate the time evolution of the system with a discrete timestep defined by the fastest rate, the rate at which cells leave the hierarchy at the topmost level, i.e., $\Delta t = 1/r_n^{\uparrow\uparrow}$. The number of microscopic events of each type, including mutations, that occur at a timestep is drawn randomly from a Poisson distribution with the appropriate mean. The number of cells the tissue produces during its lifetime ($N$) is directly proportional to the number of stem cells ($N_0$), as every stem cell creates an independent cell lineage. A single simulation instance runs until a cell accumulates the critical number of driver mutations, then we consider it cancer initiation. To measure the probability of cancer initiation, one can scale up the number of events by increasing the number of stem cells at level 0. We set $N_0$ to a number where we measure the probability of cancer initiation around 50 %. We do this by pre-estimating $N_0$ with the theoretical estimation we described in the main article, and then we fine tune the value of $N_0$ by running $10-100$ instances of the simulation (depending on the runtime) to ensure that the measured probability is around 50%, then we run the desired amount of instances to determine this probability $P_{\text{cancer}}(N_0)$ with a given precision. After the simulations we can scale the measured cancer probability to an arbitrary stem cell number $N_0'$ using the relationship: $[1-P_{\text{cancer}}(N_0')]^{1/N_0'} = [1-P_{\text{cancer}}(N_0)]^{1/N_0}$.

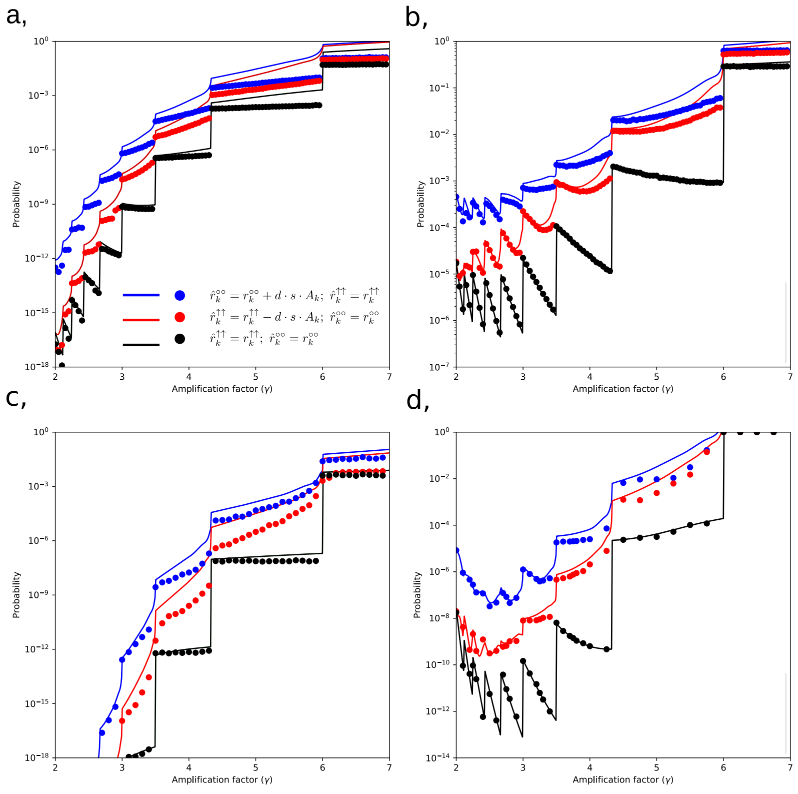


FIG. S1. **Explicit numerical validation of equation (23) for different parameter values. a)** $N/N_0 = 10^4, n=7, \mu = 4\cdot 10^{-4}$ and $s=0.1$; **b)** $N/N_0 = 10^6, n=7, \mu = 9\cdot 10^{-5}$ and $s=0.1$; **c)** $N/N_0 = 10^7, n=15, \mu = 10^{-6}$ and $s=0.1$; **d)** $N/N_0 = 10^{10}, n=15, \mu=10^{-6}$ and $s=0.1$. For each set of parameters we show three different mutational schemes: i) driver mutations increase the rate of symmetric cell division without differentiation, ii) driver mutations decrease the rate of symmetric cell division with differentiation and iii) driver mutations have no effect on rates until the critical number of mutations is accumulated. Throughout we assume $r_k^{\times} = 0$ and $N_0 = 1$.

##### B. S2.Comparison of optimal tissue organizations (Fig.3) with different lifetime incidences applied

The particular choice of threshold was based on an order of magnitude estimate of the cancer lifetime incidences, $2\%$ for blood and $4\%$ for colon, respectively (FIG.3). Although they are biologically realistic numbers, one could argue that these thresholds may seem very arbitrary. In this section, we show that our results regarding optimal tissue organization are robust because the overall structure of the plots remains the same regardless of the orders of magnitude changes in incidence threshold ($0.1\%, 1\%$ and $10\%$ respectively).

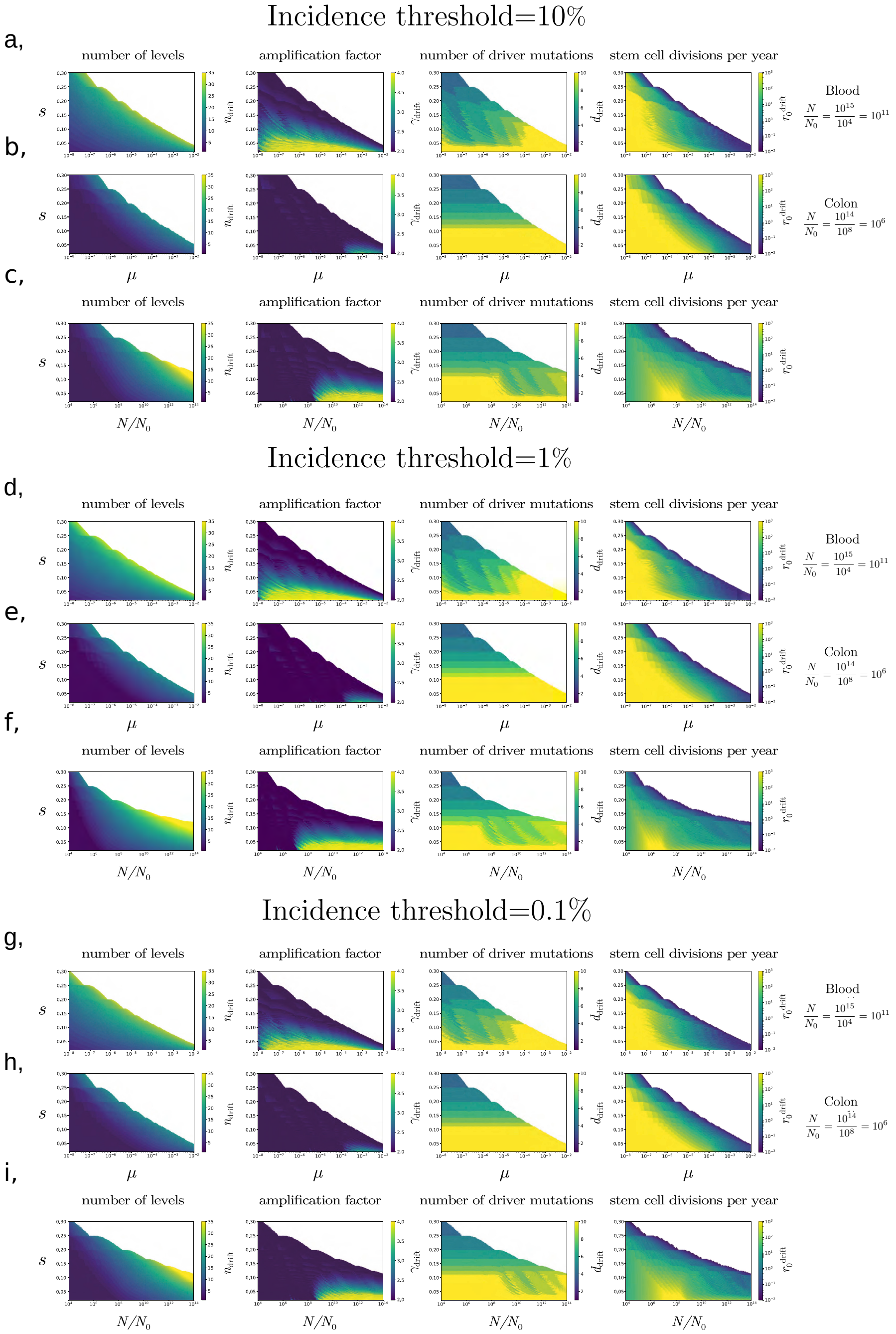


FIG. S2. **The organization of hierarchical tissues that have evolved to limit somatic evolution below thresholds between** $0.1\%$ **and** $10\%$**.** Fig.3 from the main text is reproduced with alternative values for the threshold value of cancer incidence below which selection is dominated by drift. Broad similarity can be observed for different threshold values of $0.1\%$, $1\%$ and $10\%$. Throughout we assume $r_k^{\times} = 0$.

##### C. S3.Comparison of optimal tissue organizations with neutral and drivers accumulating via only symmetric divisions.

Interestingly, optimal tissue organizations in our model are robust in the sense of different incidence thresholds and seem to be very similar if mutations can accumulate without any selective advantage (neutral mutations) and when mutations have the selective advantage (drivers). We show a comparison between optimal tissue organizations (blood and colon); in both cases, when mutations are neutral or driver mutations via symmetric divisions can accumulate in cells. We compare both in $\{N, s\}$ and $\{\mu, s\}$ heatmaps. The overall patterns are the same in both neutral and non-neutral cases.

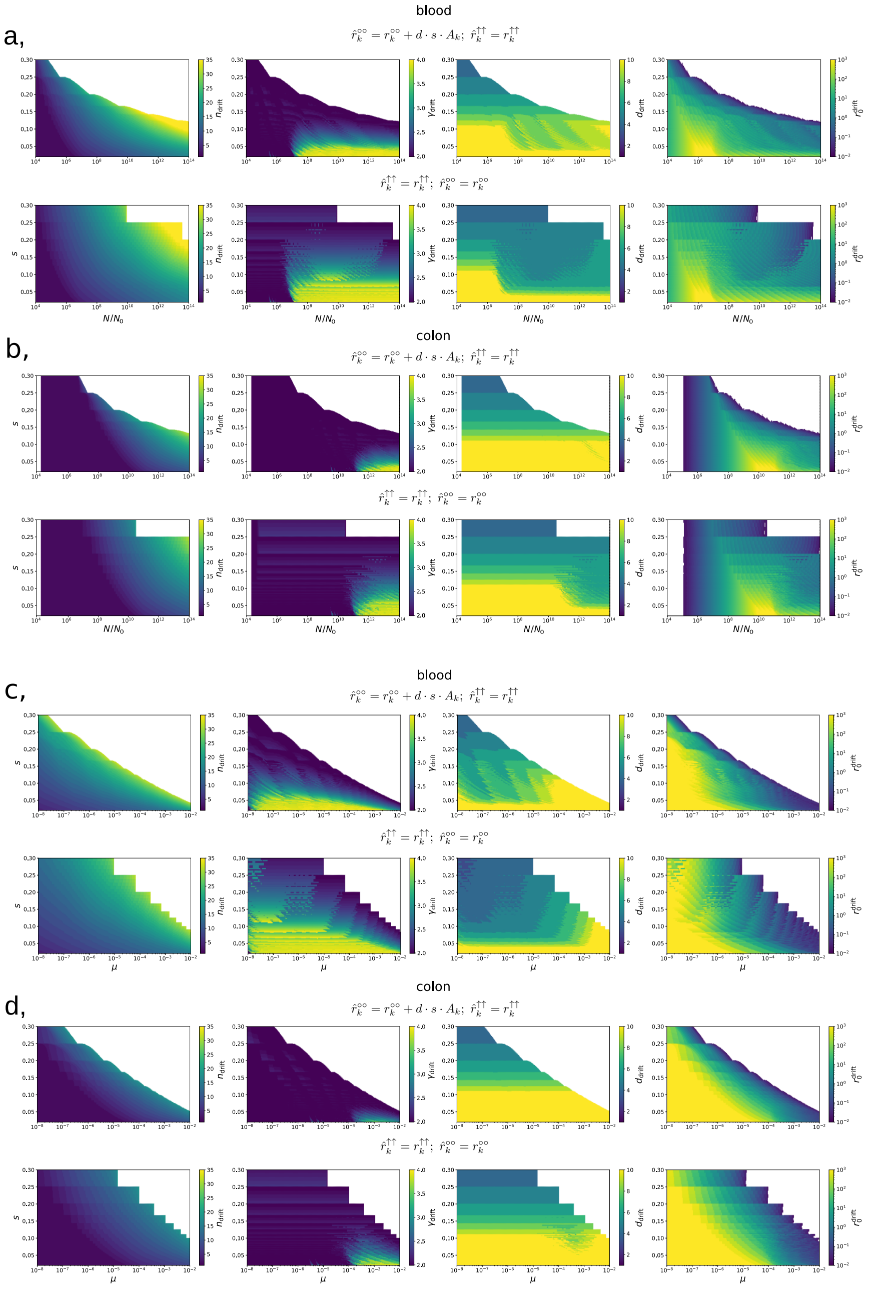


FIG. S3. **Comparison of tissue organization under two different driver mutation schemes**. We compare the organization of hierarchical tissues that have evolved to limit somatic evolution below a threshold of $1\%$ in the presence of either, **a-d** top panels, driver mutations that increase linearly the rate of symmetric cell division without differentiation (the model considered in the main text) or, **a-d** bottom panels, to driver mutations that have no effect on rates until the critical number of mutations is accumulated. Throughout we assume $r_k^{\times} = 0$.
